# Leakage-Safe Cross-Cohort Transcriptomic Survival Analysis in Pancreatic Ductal Adenocarcinoma: A Corrected Reanalysis

**DOI:** 10.1101/2025.11.14.688421

**Authors:** Mark Barsoum Markarian, Garen Vagharshag Houssigian

**Author notes:** Correspondence: Mark Barsoum Markarian. Code and results: github.com/MarkBarsoumMarkarian/RNA-harmonization-AI.

## Abstract

**Background:** Transcriptomic prognostic signatures in pancreatic ductal adenocarcinoma (PDAC) are vulnerable to endpoint misspecification, heterogeneous specimen inclusion, platform effects and information leakage. We re-audited and replaced an earlier batch-harmonized alive/dead classifier.

**Methods:** GSE71729 was audited for assay and composition but excluded from prognosis because its GEO Series Matrix does not provide an auditable patient-level survival join. Development used 176 TCGA-PAAD primary tumors with 92 deaths. A penalized Cox model was evaluated by repeated nested 5-fold cross-validation with filtering, scaling, outcome-guided screening and tuning learned inside each training partition. Cross-platform transport was tested without joint ComBat by holding out 79 survival-labelled primary tumors from GSE85916 (57 deaths), using a per-sample percentile-rank representation. Discrimination was summarized by Harrell’s concordance index (C-index) with patient-bootstrap 95% confidence intervals (CIs). The five genes highlighted in version 1 were re-examined by cohort-specific Cox models and random-effects meta-analysis.

**Results:** Internal TCGA C-index was 0.630 (95% CI 0.567-0.694). TCGA-to-GSE85916 C-index was 0.527 (0.435-0.607); reverse-direction transport was 0.576 (0.508-0.647). A fixed five-gene Cox model trained in TCGA gave external C-index 0.569 (0.476-0.657). None of the five random-effects associations survived Benjamini-Hochberg correction across the panel; heterogeneity was high (I2 74%-93%).

**Conclusions:** The corrected data support exploratory transcriptomic signal within TCGA but not a validated transportable prognostic model or validated five-gene biomarker panel. Cohort mixing after batch correction is not evidence of biological preservation or external validity. The revised open workflow provides a reproducible basis for larger, prespecified multi-cohort studies.

**Material correction notice:** This version replaces the analytical claims of version 1. GSE71729 was incorrectly described as RNA sequencing, heterogeneous specimens were treated as a prognostic validation cohort, vital status was analyzed without follow-up time or censoring, and preprocessing allowed held-out samples to influence model construction. The corrected analysis uses time-to-event outcomes, training-only preprocessing, nested validation and a survival-labelled whole-cohort test. The main conclusion changes from successful validation to modest internal discrimination without convincing cross-platform transport.

## 1. Introduction

PDAC remains a lethal malignancy, and transcriptomic profiles may contain information about tumor phenotype and outcome [1,2]. Yet prognosis modelling with high-dimensional expression data is unusually easy to overstate. Follow-up time and censoring must be represented explicitly; every data-adaptive operation must be isolated from held-out patients; and a cohort used for external testing must contain eligible patients, auditable outcomes and compatible measurements [3-7].

Version 1 of this study attempted to combine TCGA-PAAD with GSE71729 using ComBat and then classify patients as alive or dead. A forensic audit showed that the design could not support its conclusions.

GSE71729 is an Agilent microarray rather than RNA-seq and contains primary tumors, metastases, cell lines, normal pancreas and distant-site normal samples [8,9]. It was used without survival labels. Moreover, joint correction allowed the purported validation cohort to influence its own transform, and full-data preprocessing preceded out-of-bag scoring.

The purpose of this corrected reanalysis is therefore narrower and more defensible: to ask whether an expression-only survival model developed in one PDAC cohort retains discrimination in a completely held-out survival-labelled cohort measured on another platform, while keeping preprocessing and feature selection inside the appropriate training boundary. A second aim is to assess whether the five original candidates - LAMC2, DKK1, ITGB6, GPRC5A and MAL2 - remain credible as exploratory hypotheses.

**Figure 1.**
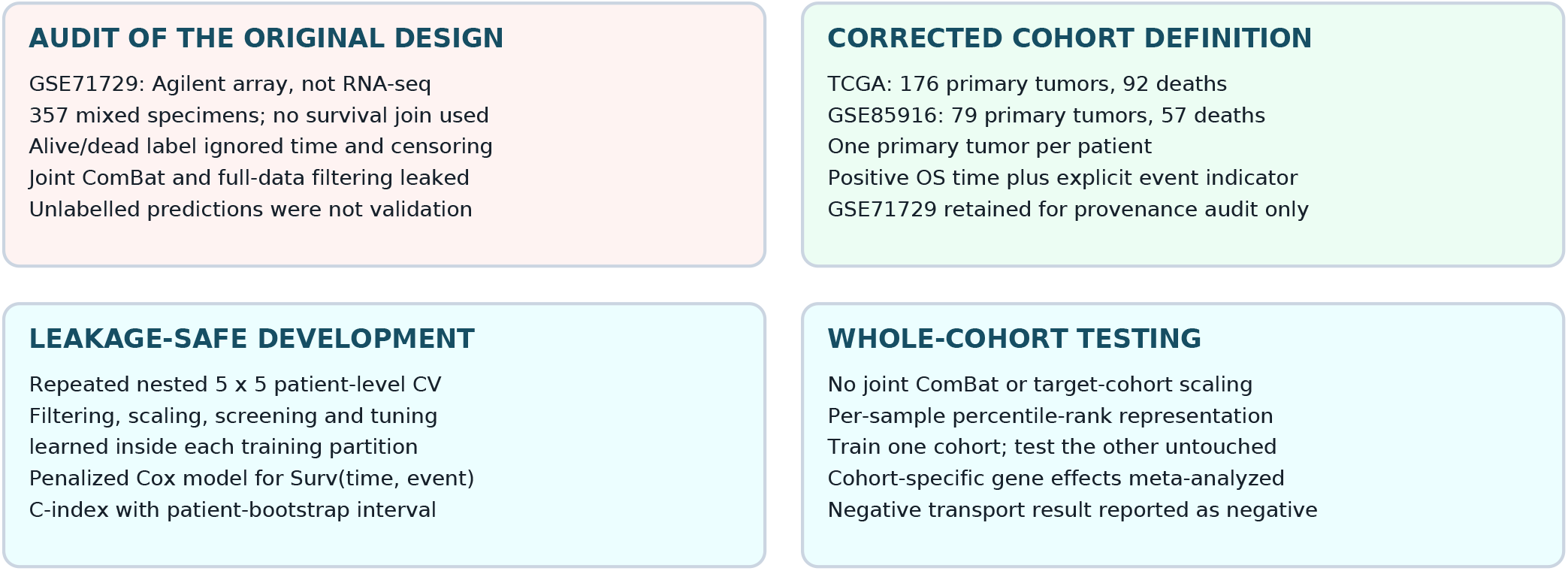
Audit and corrected study design. The upper panels summarize the defects that invalidated the original prognostic and external-validation claims. The lower panels show the replacement workflow. The target cohort never contributes expression values, survival outcomes, centering, scaling, feature selection, tuning or model coefficients to its own prediction

## 2. Materials and methods

### 2.1 Study design and correction scope

This is a retrospective corrective reanalysis of public de-identified data. The original repository state is preserved at tag v1-original-preprint; corrected scripts and machine-readable outputs are under scripts/v2 and results/v2. We followed the core prediction-model principles of defining eligible participants, outcome, unit of analysis and validation boundary before model fitting [4-7]. The reanalysis was not prospectively registered, and GSE85916 was selected during the repair; its results are therefore exploratory rather than confirmatory.

### 2.2 Cohort audit, eligibility and outcomes

TCGA-PAAD. For a fast reproducible correction, expression and clinical data were obtained from the cBioPortal PanCancer Atlas data snapshot [10,11]. Non-negative RSEM abundance estimates were transformed as log2(x+1). We retained one primary tumor per patient, required positive overall-survival time and an explicit death/censoring indicator, and matched the clinical and expression records. This yielded 176 patients and 92 deaths across 20,511 uniquely represented genes.

GSE85916. Submitter-normalized expression and sample annotations were obtained from the GEO Series Matrix. The series uses the GPL13667 Affymetrix Human Genome U219 Array and contains resected FFPE primary PDAC tumors [12,13]. Ambiguous multi-gene probes were excluded; probes mapping unambiguously to the same symbol were averaged without using outcomes. Of 80 samples, 79 had positive overall-survival time and a known event indicator, including 57 deaths, with 19,040 mapped genes.

GSE71729. The GEO record and source study identify expression profiling by GPL20769 Agilent whole-genome microarray [8,9]. We audited all 357 records by specimen title. The current Series Matrix does not provide the patient-level survival-time/event join required for this analysis; therefore, GSE71729 was not used for model development or validation.

**Table 1.** Cohorts and prognostic eligibility.

| Dataset | Assay / role | Eligible composition | Patients / events |
| --- | --- | --- | --- |
| TCGA-PAAD | Illumina RNA-seq; model development and internal validation | One primary tumor per patient; positive OS; known event | 176 / 92 |
| GSE85916 | GPL13667 Affymetrix U219; retrospective whole-cohort test | Primary resected tumors; positive OS; known event | 79 / 57 |
| GSE71729 | GPL20769 Agilent array; audit only | Mixed primary, metastatic, normal and cell-line specimens; no auditable survival join in Series Matrix | Not eligible |

OS, overall survival. Event denotes death; event=0 denotes right-censoring.

### 2.3 Internal model development

The endpoint was Surv(overall-survival time, event), not vital status alone. TCGA model development used five repeats of stratified 5-fold outer cross-validation. Within each outer training partition, a 3-fold inner loop selected the number of screened genes (5, 10 or 20) and the ridge penalty parameter (theta 0.1, 1 or 10) by mean validation C-index. Within every training split, genes were first restricted to the 1,000 largest median absolute deviations; missing values were imputed with training medians; and predictors were centered and scaled using training estimates. Features were then ranked by the absolute cross-product with martingale residuals from a training-only null Cox model. A ridge-penalized Cox model was fitted using Efron handling of ties. The same learned transform and coefficients were applied to the held-out partition.

Each patient’s standardized outer-fold risk score was averaged across the five repeats. Harrell’s C-index was calculated on these aggregated out-of-fold scores. A 95% CI was obtained from 1,000 patient-level bootstrap samples. Feature-selection stability was the proportion of the 25 outer fits selecting each gene.

### 2.4 Whole-cohort cross-platform evaluation

TCGA and GSE85916 shared 18,445 uniquely mapped gene symbols. This shared feature dictionary used platform annotation but no target-cohort values or outcomes; a future locked study should predeclare the assay panel before opening the external cohort. Each sample was independently converted to percentile ranks across its measured genes. This deployable representation avoids joint ComBat and avoids using the target cohort’s distribution for normalization. The training cohort alone supplied MAD filtering, imputation, centering, scaling, outcome-guided screening, tuning and model coefficients. We evaluated both TCGA-to-GSE85916 and GSE85916-to-TCGA directions, with the entire target cohort held out.

### 2.5 Fixed-panel and candidate-gene analyses

For the prespecified five-gene panel, sample-wise ranks were generated over the shared feature dictionary. TCGA means and standard deviations standardized the five predictors, a multivariable Cox model was fitted in TCGA, and its coefficients were applied unchanged to GSE85916. Separately, each candidate was standardized within each cohort and fitted in a univariate Cox model. Cohort log-hazard ratios were combined by a DerSimonian-Laird random-effects model. Benjamini-Hochberg adjustment was applied across the five panel-level meta-analysis tests. Because the genes were originally selected adaptively from a genome-wide analysis, this five-test adjustment does not provide genome-wide error control.

### 2.6 What was not claimed or estimated

This fast correction does not estimate clinical utility, calibration, net benefit or time-dependent AUC at prespecified horizons; it does not compare an expression score with a locked clinical baseline; and it does not establish proportional-hazards validity for each candidate. These are requirements for the next manuscript-grade analysis, not evidence supplied here. No thresholded high-risk/low-risk groups were optimized on the validation outcomes.

### 3. Results

### 3.1 The original external cohort was ineligible for the stated analysis

GSE71729 contained 145 primary tumors, 61 metastases, 17 cell lines, 46 normal-pancreas samples and 88 normal samples from distant sites. Treating all 357 as a prognostic patient cohort therefore mixed biological units and specimen types. The absence of a survival-time/event join in the Series Matrix also means that a histogram of model outputs could not evaluate discrimination, calibration or survival association.

**Table 2.** Audited GSE71729 composition. Counts reproduce the authoritative series design and source article [8,9].

| GSE71729 specimen class | n | Eligible for primary-tumor survival modelling in v2 |
| --- | --- | --- |
| Primary PDAC tumor | 145 | No: survival join absent |
| Metastasis | 61 | No: wrong specimen class |
| Cell line | 17 | No: wrong specimen class |
| Normal pancreas | 46 | No: wrong specimen class |
| Distant-site normal | 88 | No: wrong specimen class |

### 3.2 Internal discrimination was modest

Repeated nested TCGA validation produced C-index 0.630 (95% CI 0.567-0.694). This is an estimate of the complete internal pipeline, not training accuracy. It indicates some rank discrimination within TCGA but does not demonstrate clinical utility or transport to another platform.

### 3.3 Whole-cohort transport was weak or absent

When TCGA was the sole training cohort, held-out GSE85916 C-index was 0.527 (95% CI 0.435-0.607), close to chance. In the reverse direction, the model trained only on GSE85916 achieved 0.576 (0.508-0.647) in TCGA. The fixed original five-gene Cox model produced apparent TCGA training C-index 0.644 but held-out GSE85916 C-index 0.569 (0.476-0.657). The external CI includes 0.5.

**Table 3.**
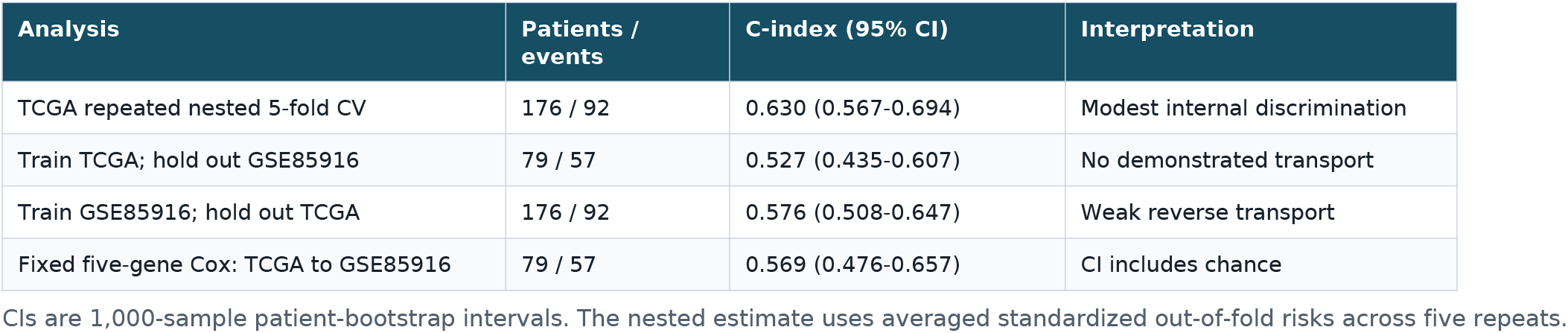
Corrected model-performance estimates.

**Figure 2.**
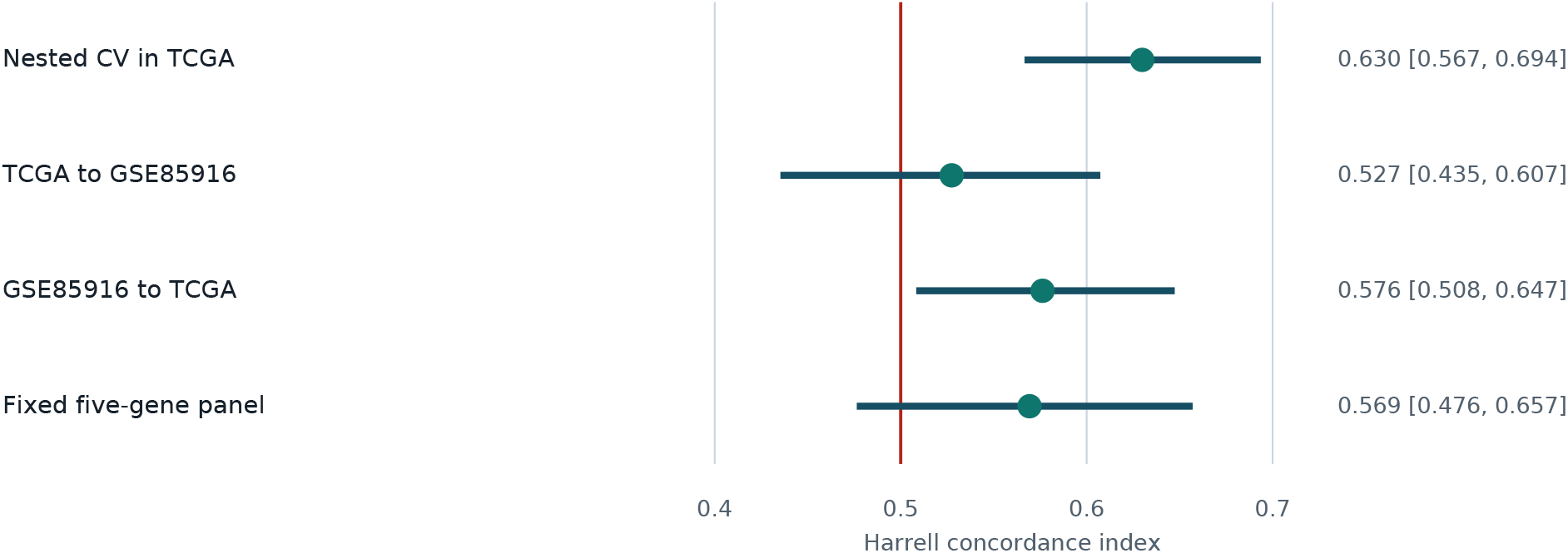
Internal and whole-cohort discrimination. Harrell concordance index Points are Harrell C-index estimates and horizontal lines are 95% patient-bootstrap CIs. The red reference is chance discrimination (0.5). The external analyses are exploratory because GSE85916 was selected during the correction.

### 3.4 Feature stability did not reproduce the original panel

Across 25 outer TCGA fits, FAM83A was selected in 25, UCA1 in 24, LY6D in 19, COL17A1 in 18 and HMGA2 in 16. DKK1 appeared in 12 fits. LAMC2, ITGB6, GPRC5A and MAL2 appeared in none. These frequencies describe internal algorithmic stability only and are not external biomarker validation.

**Table 4.**
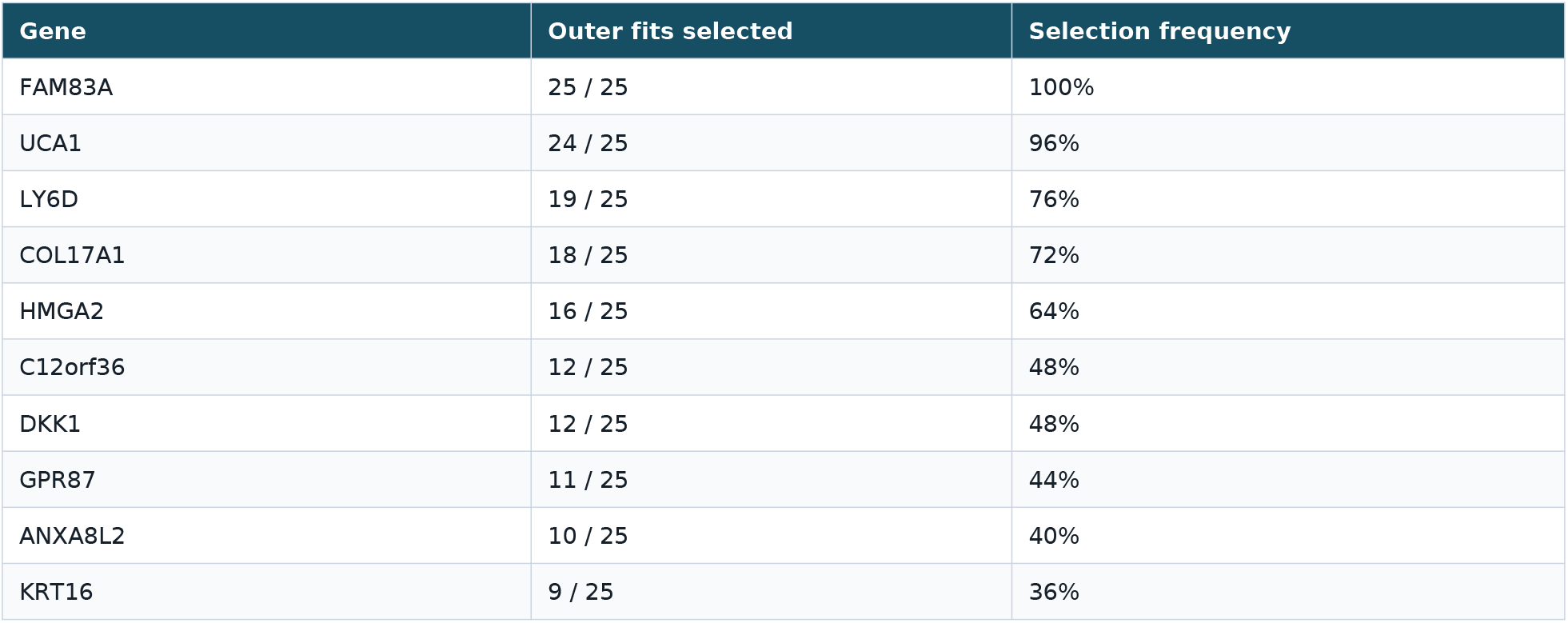
Ten most frequently selected genes in nested TCGA modelling. All selection occurred inside the corresponding outer training split.

| Gene | Outer fits selected | Selection frequency |
| --- | --- | --- |
| FAM83A | 25 / 25 | 100% |
| UCA1 | 24 / 25 | 96% |
| LY6D | 19 / 25 | 76% |
| COL17A1 | 18 / 25 | 72% |
| HMGA2 | 16 / 25 | 64% |
| C12orf36 | 12 / 25 | 48% |
| DKK1 | 12 / 25 | 48% |
| GPR87 | 11 / 25 | 44% |
| ANXA8L2 | 10 / 25 | 40% |
| KRT16 | 9 / 25 | 36% |

### 3.5 Candidate effects attenuated across cohorts

All five original candidates had adverse univariate associations in TCGA, the cohort in which they were discovered. Effects were smaller in GSE85916. In two-cohort random-effects models, LAMC2 was nominally associated with worse survival (HR 1.58 per cohort SD, 95% CI 1.04-2.42; unadjusted p=0.034), but its panel-level BH FDR was 0.170. None of the five passed BH adjustment, and I2 ranged from 73.7% to 93.0%, indicating substantial between-cohort heterogeneity.

**Figure 3.**
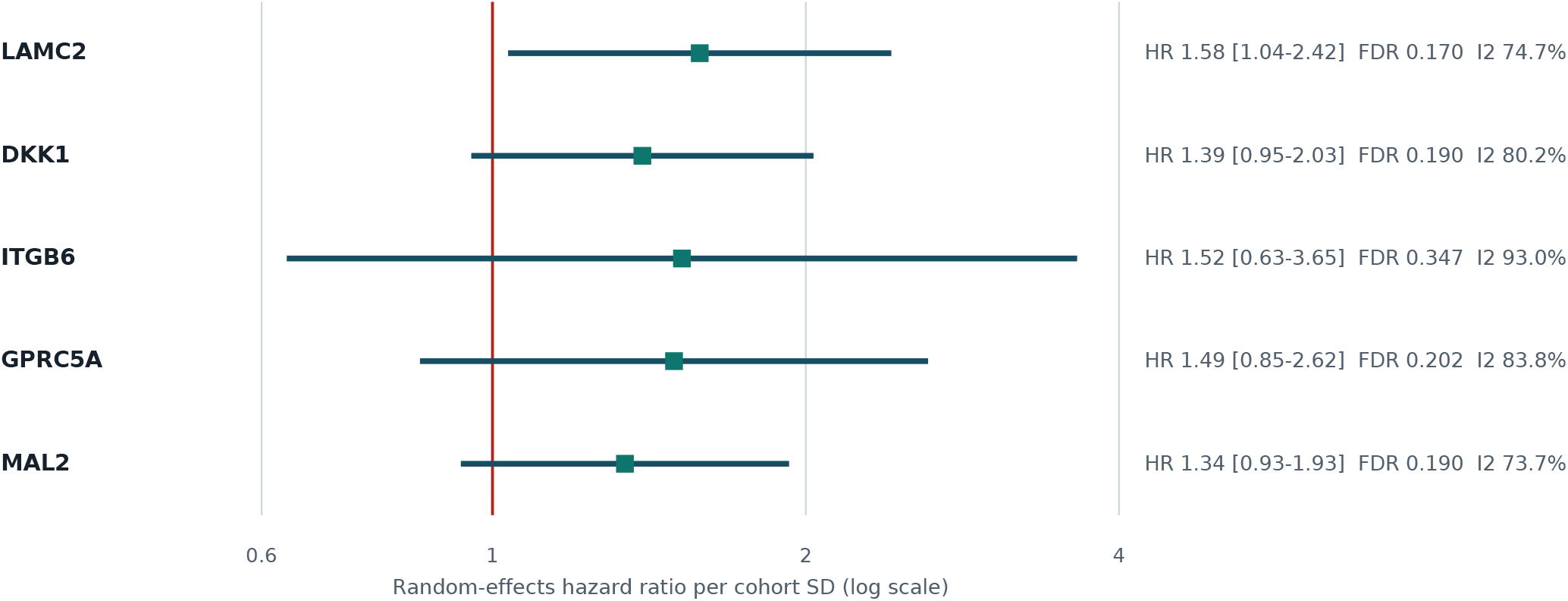
Two-cohort random-effects estimates. Random-effects hazard ratios for the five candidates. The vertical red line is HR=1. All panel-level BH FDR values exceed 0.05, and heterogeneity is high. With only two cohorts, both heterogeneity and random-effects uncertainty are imprecisely estimated.

**Table 5.** Exploratory candidate-gene survival associations.

| Gene | TCGA HR | GSE85916 HR | Random-effects HR (95% CI) | BH FDR | I2 |
| --- | --- | --- | --- | --- | --- |
| LAMC2 | 1.97 | 1.28 | 1.58 (1.04-2.42) | 0.170 | 74.7% |
| DKK1 | 1.69 | 1.15 | 1.39 (0.95-2.03) | 0.190 | 80.2% |
| ITGB6 | 2.41 | 0.99 | 1.52 (0.63-3.65) | 0.347 | 93.0% |
| GPRC5A | 2.01 | 1.13 | 1.49 (0.85-2.62) | 0.202 | 83.8% |
| MAL2 | 1.61 | 1.11 | 1.34 (0.93-1.93) | 0.190 | 73.7% |

HRs are per within-cohort standard deviation. TCGA and GSE85916 HR columns show point estimates; random-effects columns include 95% CIs. BH correction covers only five panel-level tests and not the original genome-wide selection.

### 3.6 The proposed GPRC5A paradox was not reproduced

Version 1 described lower GPRC5A expression in deceased patients as a paradox relative to experimental evidence that GPRC5A can promote PDAC cell proliferation and metastasis [14]. The corrected time-to-event analyses instead gave adverse directions in both cohorts: HR 2.01 per SD in TCGA and HR 1.13 per SD in GSE85916. The random-effects estimate was HR 1.49 (95% CI 0.85-2.62; BH FDR 0.202; I2 83.8%). Thus, the direction is no longer paradoxical, but the cross-cohort evidence is too imprecise and heterogeneous to establish a prognostic biomarker.

## 4. Discussion

### 4.1 Principal findings

This correction materially changes the scientific conclusion. Once survival time and censoring were respected, and once complete training boundaries were enforced, expression carried modest internal rank information in TCGA but did not transfer convincingly to GSE85916. The original five-gene panel likewise failed to show reliable external discrimination, and none of its random-effects associations survived even the limited five-test adjustment.

A negative transport result is informative. It shows why visually successful cohort mixing cannot substitute for outcome-labelled external evaluation. ComBat can remove cohort separation, but when cohort is confounded with platform, specimen type and disease setting, disappearance of a batch silhouette does not prove retention of prognostic biology. Joint correction also describes a transductive dataset-specific analysis rather than a model that can be applied prospectively to a new patient.

### 4.2 What remains scientifically useful

The motivating problem - reproducible PDAC transcriptomic prognosis across cohorts - remains important. The five genes also remain biologically plausible hypotheses, particularly LAMC2, DKK1 and ITGB6, whose TCGA associations were strong. Their attenuation and heterogeneity are reasons for better validation, not proof that the underlying biology is absent. Likewise, the stable TCGA features FAM83A, UCA1, LY6D, COL17A1 and HMGA2 may justify prespecified follow-up, but they were identified during resampling of a single development cohort and cannot yet be called biomarkers.

### 4.2 Why the original accuracy table was misleading

The value labelled training accuracy in version 1 was exactly one minus Random Forest out-of-bag error and should have been called OOB accuracy. The values labelled class balance (52.9% alive and 74.2% dead) were class-specific recalls; actual class counts were 85 alive and 93 dead. The raw-model recalls were 47.1% and 78.5%, but the corrected-model recalls were printed for both rows. More fundamentally, OOB predictions did not validate the full pipeline because full-data transformation and feature filtering had already occurred. The unsupported 92.6% dashboard accuracy has been removed from the corrected repository.

### 4.4 Limitations of the corrected reanalysis

The fast TCGA reconstruction uses a cBioPortal PanCancer Atlas RSEM snapshot rather than a pinned reprocessing of raw GDC STAR counts. GSE85916 uses submitter-normalized FFPE microarray data, and details of pre-analytical handling differ from TCGA. The cross-platform gene dictionary was formed from both platform annotations, and GSE85916 was selected during repair rather than locked prospectively. Only two survival cohorts were analyzed, limiting the precision of random-effects heterogeneity estimates. No clinical baseline, calibration, time-dependent AUC, decision-curve analysis, tumor-purity adjustment, treatment stratification or formal proportional-hazards diagnostics were included. C-index alone does not establish clinical value.

### 4.5 Revised analysis plan

A manuscript-grade next phase should preregister cohort eligibility, time origin, censoring rules, gene mapping and validation order; reprocess a pinned GDC release and raw array files with platform-specific methods; reserve at least one untouched primary-PDAC survival cohort; and compare locked clinical-only, expression-only and combined models. It should report C-index, prespecified time-dependent AUCs, calibration, clinical utility and uncertainty under leave-one-cohort-out validation. Candidate associations should be estimated separately by cohort, checked for proportional hazards, adjusted for key clinical covariates where consistently available, and meta-analyzed with transparent multiplicity control. Platform harmonization methods should be evaluated by predictive transport and biological retention, not mixing alone.

## 5. Conclusions

The corrected analysis does not validate the original classifier or five-gene signature. It provides evidence of modest internal transcriptomic prognostic signal in TCGA, but cross-platform whole-cohort discrimination was weak and uncertain. GSE71729 is an Agilent mixed-specimen microarray and cannot serve as the RNA-seq external cohort described in version 1. The useful outcome of this work is a transparent, open, leakage-aware workflow and a set of clearly labelled hypotheses for a larger prespecified multi-cohort study.

## 6. Transparent correction summary

| Version 1 statement or practice | Correction in version 2 |
| --- | --- |
| GSE71729 described as Illumina RNA-seq | It is GPL20769 Agilent microarray [8,9]. |
| All 357 GSE71729 samples treated as validation patients | Composition audited; mixed specimens excluded; no survival-labelled validation performed. |
| Alive/dead classification | Overall-survival time plus death/censoring indicator in Cox models. |
| Joint ComBat of development and target cohorts | No joint ComBat; per-sample rank representation for cross-platform testing. |
| Full-data filtering before OOB scoring | Filtering, imputation, scaling, screening, tuning and fitting occur inside training partitions. |
| Prediction histogram called external validation | Whole-cohort C-index with 95% CI in survival-labelled GSE85916. |
| Five validated biomarkers | Five exploratory hypotheses; none passes panel-level BH meta-analysis; high heterogeneity. |
| GPRC5A paradox | Not reproduced by corrected time-to-event analysis; direction adverse but externally uncertain. |
| Clinical/generalizable framework | Retrospective methodological correction; no clinical-use claim. |

## Acknowledgments

The authors thank the TCGA Research Network, cBioPortal Datahub maintainers and GEO data contributors for making de-identified genomic and clinical data publicly available.

## Ethics

This study used only public de-identified data and did not involve new participant recruitment or intervention.

## Author contributions

M.B.M. conceived the study, performed the computational reanalysis and drafted the corrected manuscript. G.V.H. performed a general audit, sanity checks and manuscript review. Both authors reviewed and approved the corrected manuscript.

## Data and code availability

Corrected code, input manifests, sample flows, out-of-fold predictions, model-performance tables, feature-selection frequencies and candidate estimates are available https://github.com/MarkBarsoumMarkarian/RNA-harmonization-AI. The exact repository correction is commit f42636c4782787b9ee1712782af16f3b9fa9ef19; the original state is preserved at tag v1-original-preprint. Public source data are available from TCGA/cBioPortal [10,11], GSE85916 [12] and GSE71729 [8]. Input URLs, byte sizes and MD5 hashes are included in the repository.

